# Identifying Longevity Associated Genes by Integrating Gene Expression and Curated Annotations

**DOI:** 10.1101/2020.01.31.929232

**Authors:** F. William Townes, Kareem Carr, Jeffrey W. Miller

**Affiliations:** Department of Computer Science, Princeton University, Princeton, NJ; Department of Biostatistics, Harvard T.H. Chan School of Public Health, Boston, MA

## Abstract

Aging is a complex process with poorly understood genetic mechanisms. Recent studies have sought to classify genes as pro-longevity or anti-longevity using a variety of machine learning algorithms. However, it is not clear which types of features are best for optimizing classification performance and which algorithms are best suited to this task. Further, performance assessments based on held-out test data are lacking. We systematically compare five popular classification algorithms using gene ontology and gene expression datasets as features to predict the pro-longevity versus anti-longevity status of genes for two model organisms (*C. elegans* and *S. cerevisiae*) using the GenAge database as ground truth. We find that elastic net penalized logistic regression performs particularly well at this task. Using elastic net, we make novel predictions of pro- and anti-longevity genes that are not currently in the GenAge database.

## 1 Introduction

Identifying the genetic and molecular basis of aging is a longstanding goal in medical science (Johnson & Lithgow, 1992; Remolina, Chang, Leips, Nuzhdin, & Hughes, 2012). Many studies have investigated whether individual genes are pro-longevity or anti-longevity on a case-by-case basis (Ailion, Inoue, Weaver, Holdcraft, & Thomas, 1999). Typically, an intervention such as a knockout/knockdown or overexpression is applied to a small number of genes in a model organism such as nematode worm (*Caenorhabditis elegans*) or yeast (*Saccharomyces cerevisiae*) followed by quantification of lifespan. A gene is considered *pro-longevity* if its expression is directly related to lifespan — for instance, if overexpression increases lifespan or underexpression decreases lifespan (Tacutu et al., 2018). Conversely, a gene is considered *anti-longevity* if its expression is inversely related to lifespan. Meanwhile, many genes do not fall clearly into either category, for instance, a gene might have no discernable effect on lifespan. The GenAge database (Tacutu et al., 2018) contains a catalogue of putative pro- and anti-longevity genes based on current evidence.

Pro/anti-longevity genes can be identified by intervening on individual genes, but this is slow and expensive. Alternatively, a common technique is to randomly knock out or disrupt many genes in a population of organisms, screen for the longest living individuals, and then determine which genes were disrupted in these individuals. This screening technique can rapidly identify anti-longevity genes, but systematically identifying pro-longevity genes is less straightforward. Indeed, among the small number of genes annotated as having some impact on longevity in worms and yeast, there are considerably more anti-longevity genes than pro-longevity genes.

To prioritize which genes to investigate and speed up the discovery process, recent studies have sought to computationally predict the effect of gene interventions on aging, using annotations like Gene Ontology (GO) terms (Gene Ontology Consortium, 2019) as predictors. A survey of such efforts is provided by Fabris, de Magalhães, and Freitas (2017). However, these recent studies suffer from several limitations. First, annotations like GO may be biased by the scope of the existing literature (Haynes, Tomczak, & Khatri, 2018). Second, it is difficult to compare results across studies since there is a lack of consistency in the choice of algorithms, feature sets, and predictive target/outcome. Finally, most recent studies do not report predictive performance on a held-out test dataset, leading to possible overestimation of performance.

We address these gaps by systematically assessing the performance of five popular machine learning algorithms on the task of predicting the pro-versus anti-longevity status of genes in *S. cerevisiae* and *C. elegans*. We use a consistent outcome in all comparisons based on GenAge annotations (Tacutu et al., 2018). We compare the efficacy of GO terms versus gene expression profiles as feature sets for prediction. Further, we predict possible pro/anti-longevity genes that are not currently annotated in GenAge to suggest directions for future experimental studies.

## 2 Results

### 2.1 Data sources and algorithms

We compare the performance of five machine learning classification algorithms: elastic net penalized logistic regression (pglm) (Friedman, Hastie, & Tibshirani, 2010), support vector machine with radial basis function (svm) (Karatzoglou, Smola, Hornik, & Zeileis, 2004), gradient boosted trees (xgb) (T. Chen et al., 2019), naive Bayes (nb) (Majka, 2019), and k-nearest neighbors (knn) (Schliep & Hechenbichler, 2016).

We define the outcome (that is, the target of prediction) to be the pro-versus anti-longevity annotation of individual genes from GenAge. After data cleaning, we identified 398 yeast genes and 848 worm genes with unambiguous annotations. Of these, the majority were labeled as anti-longevity (347 for yeast and 565 for worm). For validation and comparison, in yeast, we also consider replicative lifespan (RLS) outcome data for a comprehensive set of 4,698 single-gene deletions (McCormick et al., 2015). (In yeast, it is more common to use replicative lifespan rather than chronological lifespan to study aging (McCormick et al., 2015)).

As features for prediction, we consider using GO terms (Gene Ontology Consortium, 2019) and ARCHS4 gene expression profiles (Lachmann et al., 2018) for both yeast and worm. For yeast only, we also consider using the Deleteome dataset (Kemmeren et al., 2014), which contains gene expression profiles for nearly 1500 single-gene deletions. For worm only, we also consider using the Worm Cell Atlas dataset (Cao et al., 2017), which contains gene expression profiles for around 50,000 cells. We write GXP to signify Deleteome and Worm Cell Atlas for worm and yeast, respectively. Altogether, we compare the performance of five feature sets for each species: (1) ARCHS4 alone, (2) GO alone, (3) GXP alone (Deleteome for yeast, Worm Cell Atlas for worm), (4) GO combined with ARCHS4, and (5) GO combined with GXP. Normalization, filtering, and other preprocessing steps are described in the Methods section.

To predict whether a particular gene *g* is pro- or anti-longevity, we construct features in the following manner. Each GO term is considered a separate binary feature taking a value of one if gene *g* is annotated to the term and zero otherwise. For the ARCHS4, Deleteome, and Worm Cell Atlas data each experimental condition (e.g., a perturbation or tissue sample) is considered a feature and its value is given by the expression of gene *g* under that condition. Note that this is the transpose of how gene expression data are usually investigated. However, by treating experimental conditions as features and genes as observations, this allows us to exploit arbitrary gene expression data for gene *g*, not just data from when *g* is perturbed.

### 2.2 Comparative performance of algorithms and feature sets

To assess predictive performance, we use the following cross-validation scheme. For each of the two species, we split the GenAge-annotated genes into five cross-validation folds, and then for each combination of fold, algorithm, and feature set, we compute the area under the receiver-operator curve (AUC). Thus, in total, we compute 2 × 5 × 5 × 5 = 250 AUC values, 50 for each algorithm (Figures S1,S2).

To summarize the relative performance of the five algorithms, Figure 1 shows how frequently algorithm *a* has higher AUC than algorithm *b* for each pair *a, b*. More precisely, for each pair of algorithms, Figure 1 shows the fraction of times algorithm *a* has higher AUC than algorithm *b* across the 50 combinations of species, fold, and feature set. The pglm and svm algorithms consistently outperform the others in terms of AUC. The ranking of algorithms is unchanged when compared using only yeast data. Using only worm data, svm slightly outperforms pglm (0.52 instead of 0.46 in Figure 1), and knn slightly outperforms nb (0.56 instead of 0.34 in Figure 1).

**Figure 1:**
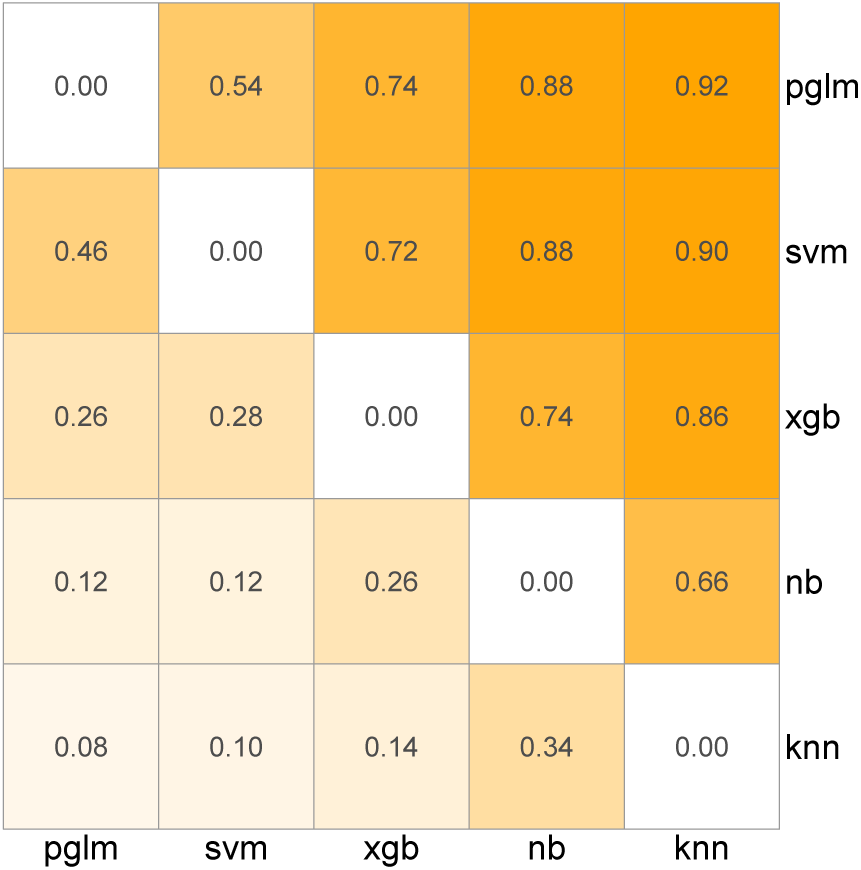
Ranking machine learning algorithms based on AUC. Numeric values indicate the fraction of times the row algorithm has higher classification performance than the column algorithm. pglm: elastic net penalized logistic regression, svm: support vector machine with radial basis function, xgb: gradient boosted trees, nb: naive Bayes, knn: k-nearest neighbors.

To compare the relative performance of the five different feature sets, Figure 2 shows boxplots of the AUC values over the five cross-validation folds, stratified by species, algorithm, and feature set. For visual clarity, here we only show the results for pglm and svm (the two best algorithms); see Figure S2 for the other algorithms. Generally speaking, using GO terms yields better predictions than gene expression features alone (ARCHS4 or GXP). However, combining GO with gene expression (GO+ARCHS4 or GO+GXP) tends to increase AUC performance compared to GO alone.

**Figure 2:**
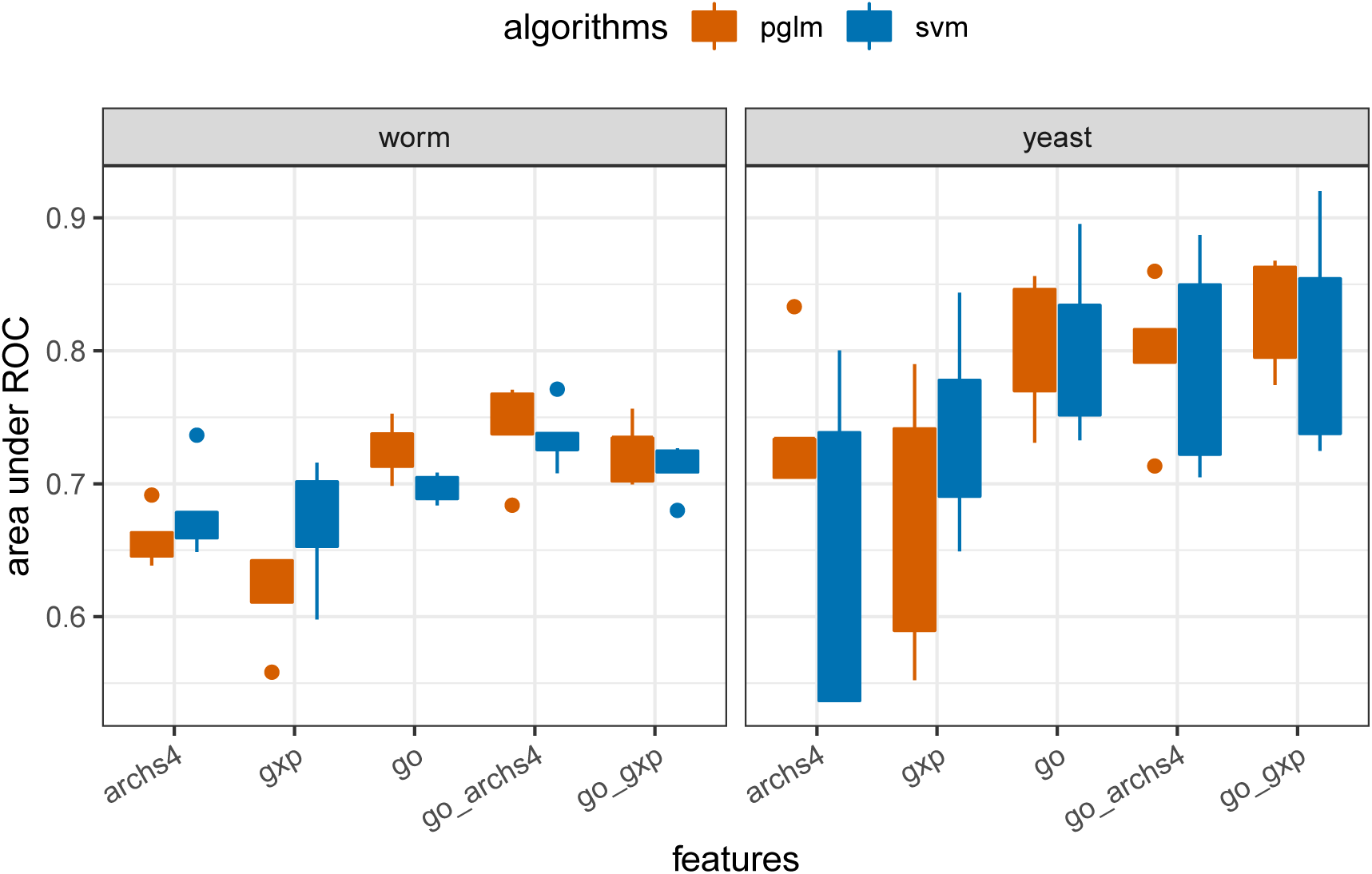
Combining gene expression (archs4, gxp) with gene ontology (GO) features yields improved classification performance in terms of AUC. pglm: elastic net penalized logistic regression, svm: support vector machine with radial basis function. An AUC value of 1 indicates perfect classification, whereas an AUC of 0.5 signifies performance no better than random.

Comparing gene expression feature sets, the ARCHS4 features give better performance than GXP (Worm Cell Atlas) for worms, but for yeast, GXP (Deleteome) is superior to ARCHS4. This could be simply due to the fact that the number of features in the worm ARCHS4 data is much larger than in the Worm Cell Atlas data. Alternatively, it could be due to the greater variation in experimental conditions across Deleteome features (which covers a comprehensive set of gene knockouts) compared to Worm Cell Atlas features (which consists of expression profiles of different cell types in normal worms).

Overall, for worms, pglm with GO+ARCHS4 features yields the best performance, whereas for yeast, pglm with GO+GXP is best (Figure 3).

**Figure 3:**
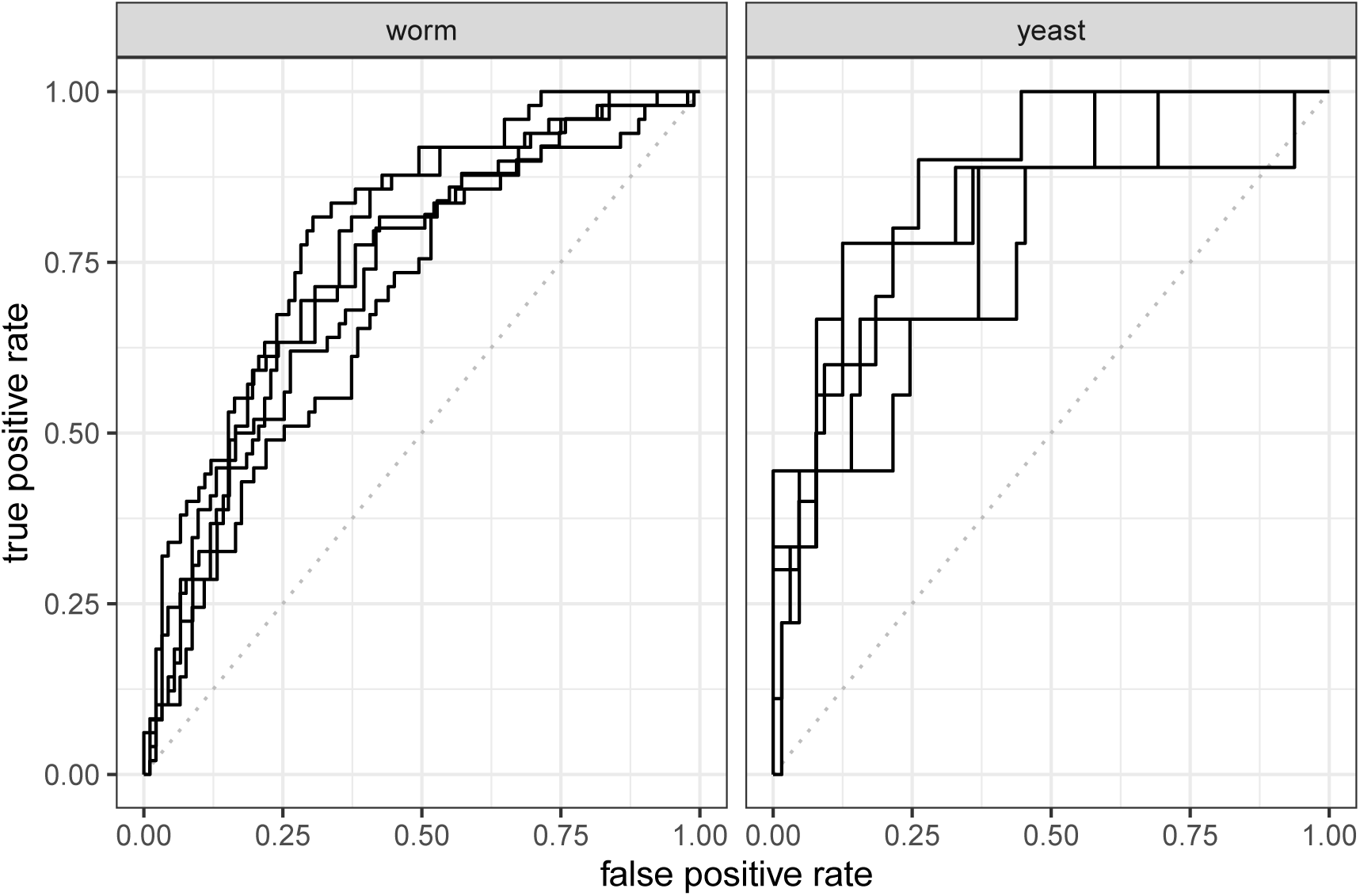
Receiver operator curves (ROC) for the best performing algorithm (pglm: elastic net penalized logistic regression) with the best performing feature sets (GO+GXP for yeast and GO+ARCHS4 for worm). Each curve represents predictive performance on the held-out data from a single cross validation fold. The diagonal gray dotted line indicates the theoretical performance of an untrained random classifier as a baseline.

### 2.3 Novel predictions of pro/anti-longevity genes

Given the encouraging performance of pglm for predicting pro/anti-longevity genes in GenAge, we applied the algorithm to make novel predictions of pro/anti-longevity genes in *C. elegans* (worm) and *S. cerevisiae* (yeast). To do this, for each species separately, we retrained a pglm model on the full GenAge database, using the combined GO terms plus ARCHS4 gene expression as features (see the Methods section for details on hyperparameter selection). We then used the trained model to generate a predictive score for the pro/anti-longevity effect of each gene not in the GenAge database. Specifically, the predictive score is defined to be the probability that the gene is pro-longevity under the trained model. A score close to 1 indicates that the gene is predicted to be pro-longevity, whereas a score close to 0 indicates that the gene is predicted to be anti-longevity. An intermediate score indicates a gene with unclear pro- or anti-longevity status. Table 1 shows the unannotated genes with the highest confidence levels of being pro- and anti-longevity for worm and yeast, respectively. These genes do not significantly overlap with predictions from the pglm model trained using only GO terms as features (Tables S2-S5), suggesting that these predictions are not simply recapitulating the known biology represented in the GO terms.

**Table 1:**
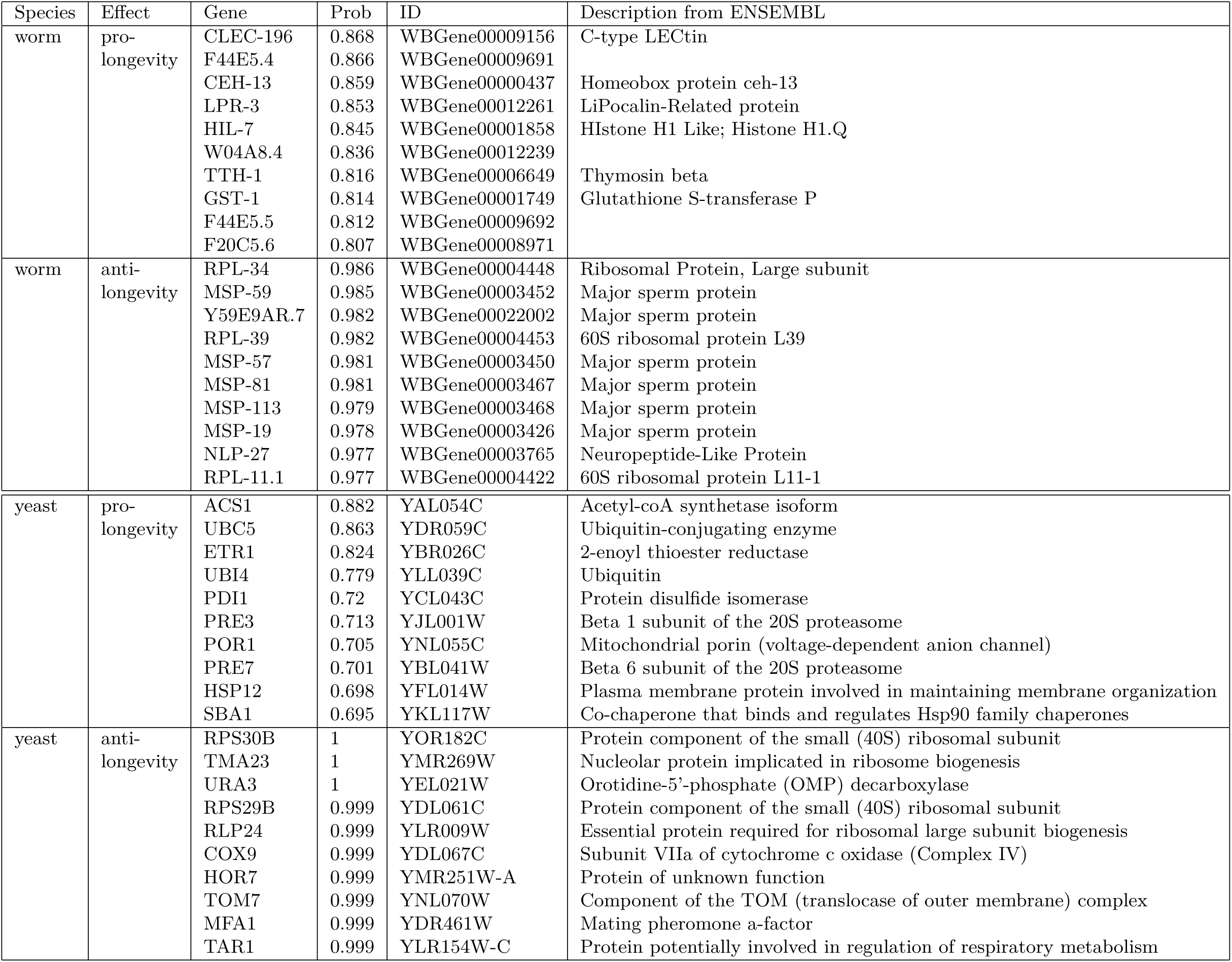
Top pro-longevity and anti-longevity genes not in GenAge predicted using GO terms and ARCHS4 gene expression for worm and yeast with the pglm (GLM-Net) algorithm.

To assess the accuracy of the predictions, we looked at the literature to see if there is experimental evidence of pro/anti-longevity effects for these genes. Based on the existing experimental evidence, we find that the model predictions are remarkably good. It turns out that—even though they are not in GenAge yet—there is experimental evidence for the pro/anti-longevity status of most of the predicted genes.

#### 2.3.1 Predicted pro-longevity worm genes

For many of the predicted pro-longevity genes in Table 1, there already exists direct experimental evidence of pro-longevity status. Note that this evidence was not used in making the predictions, implying that the model is producing reliable out-of-sample predictions. We discuss what is known about the top 10 predicted pro-longevity genes: CLEC-196, F44E5.4, CEH-13, LPR-3, HIL-7, W04A8.4, TTH-1, GST-1, F44E5.5, and F20C5.6.

F44E5.4 and F44E5.5 encode members of the *hsp70* family of heat shock proteins. The heat shock response is well-known to have strong pro-longevity effects in *C. elegans*. Indeed, knocking in extra copies of *hsp70* extends lifespan (Yokoyama et al., 2002) and knocking down *hsp70* via RNAi decreases lifespan and leads to rapid aging phenotypes (Kimura, Tanaka, Nakamura, Takano, & Ohkuma, 2007). GST-1 (Glutathione S-transferase P) is also involved in stress response—particularly, immune response—and GSTs are well-known to be pro-longevity. Overexpression (underexpression) of GSTs has been found to increase (decrease, respectively) lifespan and stress resistance (Ayyadevara et al., 2007, 2005). W04A8.4 is an uncharacterized protein that is involved in the pro-longevity effect of metformin on *C. elegans* (Wu et al., 2016); specifically, knockdown of W04A8.4 leads to metformin resistance. This is intriguing, since metformin treatment has been shown to promote health and extend lifespan in many organisms. Homeobox protein CEH-13 exhibits pro-longevity characteristics based on experimental evidence — specifically, a *ceh-13* mutant strain has decreased lifespan compared to wildtype controls (Tihanyi et al., 2010). LPR-3 (LiPocalin-Related protein) is known to be involved in nematode worm locomotion, and appears to mediate the longevity-inducing effect of *daf-7* mutation (Hyun, Kim, Dumur, Schroeder, & You, 2016); additionally, expression of *lpr-3* is increased in worms fed with *rBmα*TX14, an *α*-neurotoxin that increases worm lifespan (L. Chen et al., 2016).

For the remainder of the genes in Table 1, there is suggestive experimental evidence of pro-longevity status based on associations. C-type Lectin *clec-196* expression increases and lifespan increases when *hsb-1* is knocked out (Sural, Lu, Jung, & Hsu, 2019). Also, *clec-196* is directly adjacent to *hsp-1* on chromosome IV, suggesting possible co-involvement, and *hsp-1* (heat shock protein) is well-known to be pro-longevity. HIL-7 (Histone H1 Like) gene expression may be associated with Ethosuximide treatment, a drug that increases worm lifespan and affects DAF-16/FOXO target gene expression (X. Chen et al., 2015). TTH-1 (Thymosin beta) is significantly increased in *daf-2* mutants, which are very long-lived, suggesting possible pro-longevity status by association (Narayan et al., 2016). F20C5.6 is affected by the well-known longevity genes *clk-1* and *sir-2*.*1*, as well as by treatment with 1-methylnicotinamide and rotenone, which are well-known for increasing worm lifespan.

This validating evidence from the literature indicates that the model predictions are surprisingly accurate. The predicted pro-longevity genes CLEC-196, HIL-7, TTH-1, and F20C5.6 are candidates for further experimental exploration.

#### 2.3.2 Predicted anti-longevity worm genes

Similarly to the predicted pro-longevity genes, there exists experimental evidence of anti-longevity status of most of the predicted anti-longevity genes in Table 1. We discuss what is known about the top 10 predicted anti-longevity genes: MSP-59, Y59E9AR.7, RPL-39, MSP-57, MSP-81, MSP-113, MSP-19, NLP-27, and RPL-11.1.

Major sperm proteins appear to be anti-longevity based on the experimental evidence. A mutation reducing sperm production leads to significantly increased lifespan (Van Voorhies, 1992). Additionally, the expression of sperm-related genes—especially major sperm protein (MSP) genes—is decreased in adult *daf-2* mutants, providing further support for an anti-longevity role of MSP genes (Halaschek-Wiener et al., 2005). RSP-39 and RPL-11.1 are 60S ribosomal proteins. RNAi knockdown of genes encoding ribosomal proteins consistently increases lifespan in *C. elegans*, both in the case of 40S and 60S ribosomal proteins (Hansen et al., 2007). This supports the predicted anti-longevity status.

NLP-27 (Neuropeptide-Like Protein) is the only other predicted anti-longevity gene in the top 10 list. Expression of *nlp-27*, along with other *nlp* genes, is increased in long-lived *daf-2* mutants. Further, *nlp-27* expression is reduced in a short-lived *mir-71* deletion strain. This indirect evidence by association suggests a possible pro-longevity role of NLP-27—which would contradict the predicted anti-longevity—but direct over/under-expression of *nlp-27* would be needed to establish its pro/anti-longevity status.

#### 2.3.3 Predicted pro-longevity yeast genes

Table 1 lists the top 10 predicted pro-longevity yeast genes. Several of these predictions are borne out by direct experimental evidence via single-gene deletions — specifically, deletion of ACS1, ETR1, UBI4, and POR1 leads to decreased lifespan (Marek & Korona, 2013). Marek and Korona (2013) did not find a significant pro- or anti-longevity effect for UBC5, HSP12, or SBA1, and they do not report results for the remainder of the top 10 genes. However, UBC5 is a strong pro-longevity candidate, since it is involved in cellular stress response and mediates selective degradation of short-lived and abnormal proteins (Seufert & Jentsch, 1990). HSP12 (heat shock protein) is required for the lifespan-extending effect of dietary restriction in yeast (Herbert et al., 2012), validating the pro-longevity prediction. SBA1 is also a strong pro-longevity candidate, as a chaperone-binding protein that is involved in heat shock response and is required for telomere length maintenance (Fang, Fliss, Rao, & Caplan, 1998; Toogun, Zeiger, & Freeman, 2007). PRE3 and PRE7 are part of the proteasome, and it is known that increased proteasome capacity extends lifespan (Kruegel et al., 2011), providing indirect validation of their predicted pro-longevity status. PDI1 is a downstream target of the unfolded protein response (UPR), which is well-known to be pro-longevity (Patil & Walter, 2001).

#### 2.3.4 Predicted anti-longevity yeast genes

Table 1 lists the top 10 predicted anti-longevity yeast genes. As in worms, depletion of ribosomes increases lifespan (Steffen et al., 2008), validating the predictions of the ribosome-biogenesis proteins RPS30B, TMA23, RPS29B, and RLP24 as anti-longevity. HOR7 is reported to influence lifespan, but the direction of the effect may be context-dependent: HOR7 deletion increases lifespan (McCormick et al., 2015), whereas Schleit et al. (2013) find that HOR7 deletion decreases lifespan under dietary restricted conditions.

For URA3, COX9, TOM7, MFA1, and TAR1, we do not find pre-existing corroboration of the predicted anti-longevity status in the literature. TOM7 deletion has been reported to decrease chronological lifespan (Garay et al., 2014), and it does not appear to have a strong effect on replicative lifespan (Marek & Korona, 2013). TOM7 is part of the translocase of the outer mitochondrial membrane (TOM) complex, and the mitochondrial membrane is well-known to be important in yeast longevity (Jazwinski, 2005). Marek and Korona (2013) report a pro-longevity effect for COX9, contrary to the model prediction. (Except for COX9, the Marek and Korona (2013) results are inconclusive for all of the genes in Table 1.) Further investigation of URA3, COX9, TOM7, MFA1, and TAR1 might be interesting to pursue.

### 2.4 Validation on a secondary dataset

To further evaluate the predictive accuracy of the trained pglm model, we compare the model predictions to actual lifespan measurements from a non-GenAge validation dataset. For this purpose, we use a dataset of replicative lifespan for a comprehensive set of 4,698 single-gene deletions in yeast (McCormick et al., 2015). Since the McCormick et al. (2015) dataset contains lifespan measurements for deletions of many genes that do not appear in GenAge, in principle it should be well-suited as a secondary validation dataset. Using the pglm model trained on the full GenAge database for yeast with the GO+ARCHS4 feature set as predictors, we made predictions of the longevity effect of all 4,698 genes in the McCormick dataset.

First, as a sanity check, we observe that among genes in GenAge, the predicted probability of a gene being pro-longevity is clearly inversely related to the change in lifespan after deletion (Figure 4, left panel). This is not surprising since it simply means that the GenAge annotations are roughly consistent with the McCormick data, and the model was able to fit the GenAge-based training data. More interestingly, we see that the model is able to predict which genes have a larger or a smaller effect on lifespan (Figure 4, left panel). For instance, among pro-longevity genes, the genes with predicted probability near 1 do indeed tend to lead to a larger decrease in lifespan. Meanwhile, among anti-longevity genes, the genes with predicted probability near 0 do indeed tend to lead to a larger increase in lifespan. Since the training data contain no information about the magnitude of the effect on lifespan, this indicates that the model is not simply recapitulating the training data, but is indeed making generalizable predictions.

**Figure 4:**
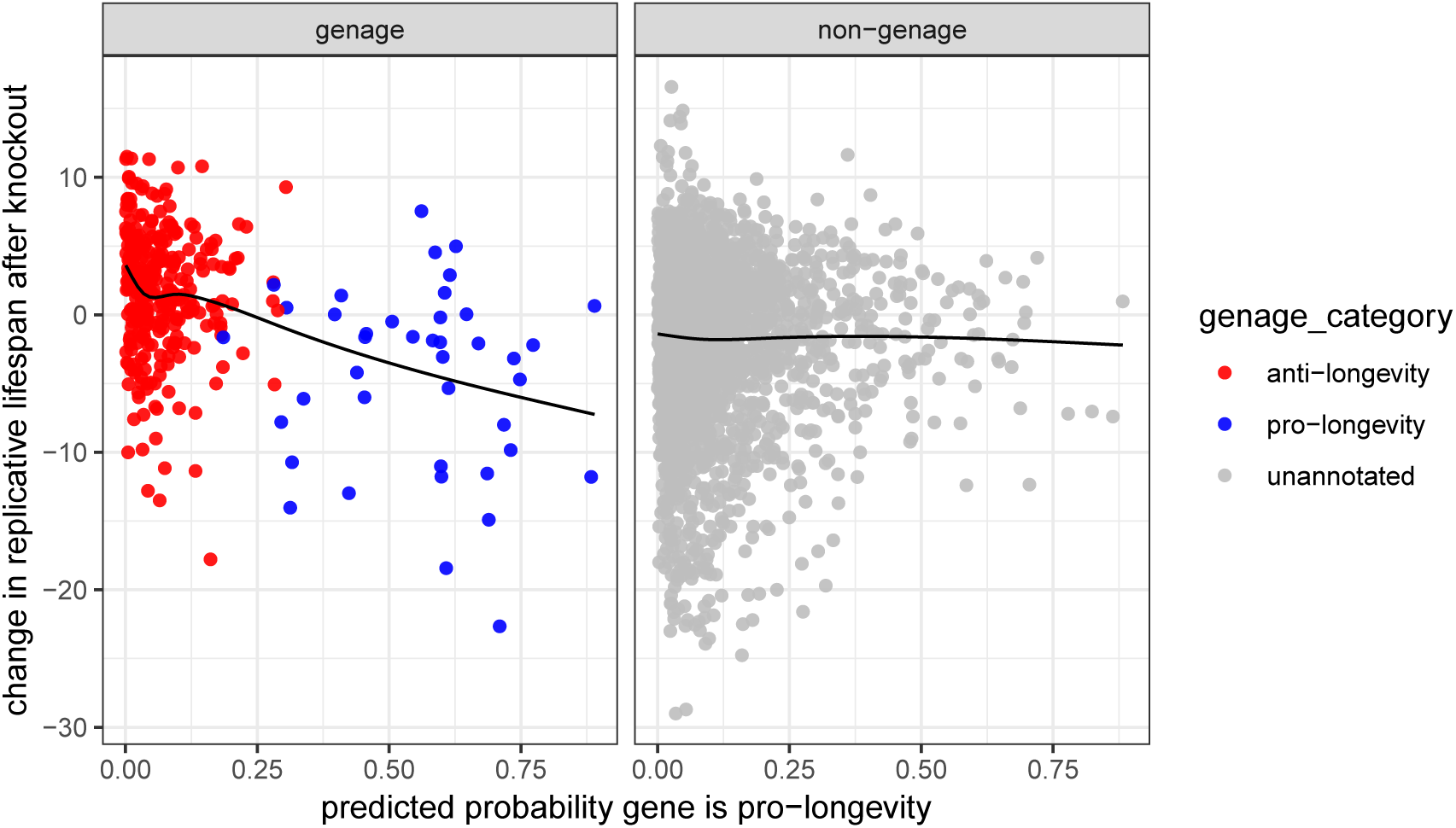
Predicted probability of a gene being pro-aging versus effect of deletion on replicative lifespan (RLS) in yeast. Probabilities are from the pglm classifier trained on the full GenAge dataset. Solid curve is a nonparametric smoother.

Next, we compare the model predictions to the lifespan data for genes outside the GenAge database. Figure 4 (right panel) shows the change in lifespan versus the predicted probability of a gene being pro-longevity, for genes in the McCormick dataset that are not in GenAge. A downward trend in this plot would indicate concordance between model predictions and the validation data. There is an extremely slight but not convincing downward trend; thus, while suggestive, this does not provide a compelling out-of-sample validation of the model predictions. Note that the pglm classifier trained on GenAge has a strong bias toward predicting genes to be anti-longevity; see Figure 4 (right panel) and Figure S3. This bias is due to class imbalance in the training data, since the majority of genes annotated in GenAge are anti-longevity. This is common when the training data are imbalanced, and can easily be addressed by selecting the classification threshold to yield appropriately balanced predictions.

The lack of concordance between the out-of-sample model predictions and the McCormick lifespan data may be attributable to the fact that for many genes, the McCormick data are not in agreement with the GenAge annotations of pro/anti-longevity. Specifically, many putatively pro-longevity genes led to large increases in lifespan when deleted, and many putatively anti-longevity genes led to large decreases in lifespan when deleted (Figure 4, left panel). It is not clear whether this discrepancy is primarily due to limitations of the GenAge database (e.g., bias and relatively small sample size) or limitations of the McCormick assay. Focusing on the latter possibility, recent studies have identified mechanisms by which disruption of a gene through knockout can activate compensatory mechanisms leading to a dramatically different phenotype than disruption of the same gene through knockdown, which reduces but does not eliminate expression (Wilkinson, 2019). If deletion of a single gene activates similar compensatory mechanisms in yeast, then this could explain the lack of concordance, since it would imply that the change in lifespan under a single-gene deletion is not necessarily related to that gene’s pro/anti-longevity status. A comprehensive assay of knockdowns (rather than deletions or knockouts) would shed light on this intriguing question.

### 2.5 Functional interpretation of model predictions

To interpret the biological basis for the model predictions in terms of functional categories, for each species we retrained the pglm model on the full GenAge dataset using only GO terms as features. We extracted the 20 most influential GO terms from the trained model by ranking the regression coefficients from largest to smallest in absolute value (Table 2). Note that in this model, the coefficient is equal to the log-odds ratio (logOR) of a gene being pro-longevity when it is annotated to a GO term versus when it is not annotated to that GO term. If a GO term has a positive logOR value, then genes annotated with that GO term are more likely to be pro-longevity under the model. Conversely, a negative logOR indicates that genes annotated with that GO term are more likely to be anti-longevity.

**Table 2:**
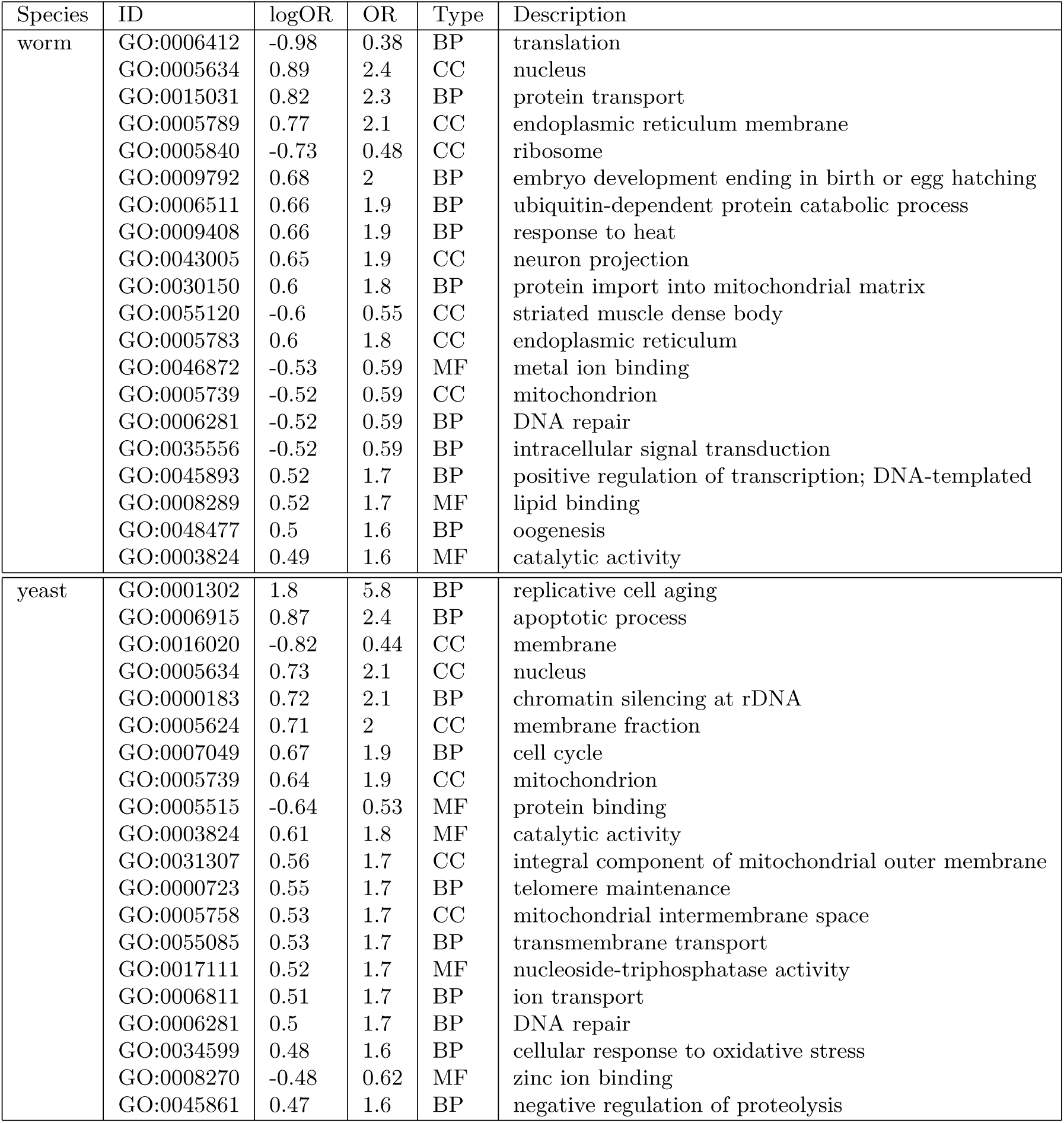
Top GO terms identified by the pglm (GLM-Net) algorithm. logOR: log-odds ratio. OR: odds ratio. Positive logOR indicates a gene annotated to that GO term is more likely to be pro-longevity. BP: biological process, CC: cellular component, MF: molecular function.

#### 2.5.1 Top GO terms for worm

The current literature supports a strong longevity effect for many of the top categories in Table 2. Translation inhibition is known to increase lifespan (Hansen et al., 2007), so a large negative coefficient for the *translation* and *ribosome* GO terms makes sense. Protein homeostasis is known to be key to longevity (Sampaio-Marques & Ludovico, 2018), so it makes sense that the model has positive coefficients for *protein transport, endoplasmic reticulum membrane*, and *endoplasmic reticulum*. Ubiquitin-mediated proteolysis is known to be important for promoting longevity, implying that a positive coefficient for *ubiquitin-dependent protein catabolic process* makes sense. Heat shock response is known to extend lifespan, and indeed, the model has a positive coefficient for *response to heat*. Activation of the mitochondrial unfolded protein response is known to promote longevity (Durieux, Wolff, & Dillin, 2011), so a positive coefficient for *protein import into mitochondrial matrix* makes sense. Mitochondria are known to be important for longevity (Sun, Youle, & Finkel, 2016), so a large coefficient for *mitochondria* makes sense; further, inhibition of mitochondrial respiration is known to extend lifespan (B Hwang, Jeong, & Lee, 2012), so a negative sign for the coefficient could make sense. Similarly, the importance of *DNA repair* makes sense, and surprisingly, in some cases, DNA repair gene knockdown increases lifespan, possibly due to compensatory biological mechanisms (Lans et al., 2013); thus, a negative coefficient is, in fact, consistent with the literature.

#### 2.5.2 Top GO terms for yeast

For yeast, Table 2 shows the top longevity-related GO terms in the model. The importance of these terms is consistent with the current literature, but the appropriate sign of the coefficient is not always clear, since the genes annotated to each GO term may have contradictory pro/anti-longevity effects and further, there may be compensatory relationships between terms due to correlated predictors.

*Replicative cell aging, apoptotic process*, and *cell cycle* obviously make sense as related to yeast aging and longevity. Mitochondrial membrane maintenance is known to be important in yeast longevity (Jazwinski, 2005), and other membranes (e.g., the vacuole membrane) may also be important (Carmona-Gutierrez, Hughes, Madeo, & Ruckenstuhl, 2016); thus, large coefficients for *mitochondrion, integral component of mitochondrial outer membrane, mitochondrial intermembrane space, membrane, membrane fraction*, and *transmembrane transport* are consistent with the literature. Depletion of ribosomes is known to increase lifespan (Steffen et al., 2008), so a negative coefficient for *chromatin silencing at rDNA* is appropriate. Telomeres are known to be important in yeast longevity (Austriaco & Guarente, 1997; Liu, Wang, Wang, & Liu, 2019), so a large coefficient for *telomere maintenance* makes sense. Longevity effects of *cellular response to oxidative stress* are corroborated in the literature (Postnikoff, Johnson, & Tyler, 2017). Finally, a negative coefficient for *zinc ion binding* is consistent with experimental evidence that zinc limitation extends chronological lifespan (Shimasaki et al., 2017).

## 3 Discussion

We systematically compared the performance of popular machine learning algorithms in classifying genes as pro- or anti-longevity using the GenAge database and combinations of gene expression and gene ontology (GO) feature sets. We identified elastic net penalized logistic regression (pglm) as the most effective classifier and made predictions for unannotated genes. The pglm model fit to GenAge data was only weakly concordant with the McCormick replicative lifespan (RLS) assay, which was based on single gene deletion strains in yeast. This discrepancy could be due to compensatory mechanisms which are known to mitigate the effects of knockouts (Wilkinson, 2019). We suggest future comprehensive longevity assays should consider knockdowns instead of deletions and knockouts. Furthermore, there is a need for increased focus on pro-longevity genes rather than anti-longevity genes, since the latter are much more common in the GenAge database. We offer our predictive probability scores as one possible tool to prioritize future experimental studies which can validate individual genes as pro-longevity mechanistically. We encourage computational researchers to use metrics such as area under receiver-operator curve (AUC) on held-out data from standard databases such as GenAge to assess classification performance and facilitate comparisons across studies. Finally, our approach of combining feature sets to improve predictive performance is generalizable in principle to a wider variety of model organisms as more annotations and datasets become available over time.

## 4 Experimental Procedures

Binary pro/anti-longevity annotations were accessed from the GenAge model organisms database (Tacutu et al., 2018), currently available at http://genomics.senescence.info/genes. We used the subset of genes for yeast and worm, and we excluded ambiguous annotations (e.g., if GenAge lists two studies for a gene, one finding it to be pro-longevity and the other finding it to be anti-longevity). GO annotations for all genes were downloaded from the BioMart ENSEMBL database using the biomaRt package in Bioconductor. For both species, gene expression data in the form of RNA-Seq read counts were obtained from the ARCHS4 database (Lachmann et al., 2018), currently available at https://amp.pharm.mssm.edu/archs4/archs4zoo.html. For yeast only, we acquired the Deleteome gene expression microarray dataset (Kemmeren et al., 2014), currently available at http://deleteome.holstegelab.nl. For worm only, we obtained gene expression data from the single-cell RNA-Seq Worm Cell Atlas (Cao et al., 2017), currently available at http://atlas.gs.washington.edu/worm-rna. We reduced the dimensionality of the Worm Cell Atlas data by summing the unique molecular identifier (UMI) counts across all cells within the same tissue, so that each feature is a “pseudobulk” tissue rather than a single cell.

Replicative lifespans (RLS) for 4,698 single-gene deletion yeast strains were obtained from the authors of McCormick et al. (2015). Perturbation genotypes with percent_change greater than 30 and set_lifespan_count less than or equal to 5 were excluded based on the authors’ recommendations. We merged results for the same genotype across replicate experiments in the following way. The outcome for each genotype in a single replicate was quantified as the mean of RLS in the perturbation group minus the mean of RLS in the control group. To obtain a single value for the genotype across all replicates, we then computed a weighted average of the outcome values from each replicate, where the weights corresponded to the sample sizes in each group. This ensured that replicates with more observations contributed more to the final value. We refer to this as the McCormick dataset.

All gene expression measurements were normalized to account for sample-specific biases. Specifically, the Deleteome data were already normalized, the ARCHS4 read counts were converted to transcripts-per-million (TPM), and the Worm Cell Atlas UMIs were converted to counts-per-million (CPM). The normalized counts were then log transformed with a pseudocount of one. For Deleteome, genes that were variable in controls and non-responsive mutants were excluded, since these data were likely to contain mostly noise. For each species, we used the subset of genes with no missing values across all feature types (GO features and the two sources of gene expression features), resulting in 703 worm genes (246 pro-longevity, 457 anti-longevity) and 368 yeast genes (46 pro-longevity, 322 anti-longevity). Features with no variation across the included genes were discarded. For yeast, the number of retained features was 3268, 700, and 1390 for ARCHS4, Deleteome, and GO terms, respectively. For worms, the number of features was 2935, 270, and 2051 for ARCHS4, Worm Cell Atlas, and GO terms, respectively. All gene expression features were centered and scaled to have mean zero and standard deviation 0.5 as suggested by Gelman (2008), while binary features (GO) were not centered and scaled. The five sets of features considered for each species were (1) ARCHS4 alone, (2) GO alone, (3) GXP alone (Deleteome for yeast, Worm Cell Atlas for worm), (4) GO combined with ARCHS4, and (5) GO combined with GXP.

To assess predictive performance of different combinations of feature sets, each dataset (consisting of the binary GenAge outcome for a single species matched with one of the five feature sets) was split into 5 external cross-validation (CV) folds. Within each fold, machine learning classifiers were fit to the training data using the caret R package (Kuhn et al., 2019). The same partitioning of the data was preserved across algorithm runs to ensure identical training and test conditions. The algorithms used were k-nearest neighbors (knn), naive Bayes (nb), gradient boosted trees (xgb), support vector machine with radial basis function (svm), and logistic regression with elastic net penalty (pglm). Hyperparameters (Table S1) were selected by grid search using repeated 10-fold internal CV with two repeats within each training fold using the Kappa criterion. Note that this means each algorithm could potentially use different hyperparameter values across the five external CV folds. For all algorithms except naive Bayes, the grid consisted of default caret values. For naive Bayes, the Laplace correction was set to zero, kernel smoothing was always used, and the adjustment to the probabilities was chosen between 0.5 and 1.0. Additionally, for naive Bayes only, many features with near-zero variance caused numerical instabilities and were excluded. Having chosen a final set of hyperparameters for each training fold, the predicted probabilities were computed for the held-out test data and the area under the receiver-operator curve (AUC) was computed to quantify prediction performance (discrimination). An AUC value of 1 indicates perfect classification performance, whereas an AUC of 0.5 signifies performance no better than random, or simply always predicting the majority class.

For the results in Sections 2.3 and 2.4, the best-performing algorithm (pglm) was retrained on all of the GenAge data for each species with the combined GO plus ARCHS4 feature set. The hyperparameter grid was expanded to 21 alpha values (evenly spaced between zero and one, inclusive), and 97 automatically selected lambda values using five-fold CV. For worm, the optimal alpha was 0.05 (close to an L2 ridge penalty). For yeast, the optimal alpha was 0.5 (an even mix between ridge and the L1 lasso penalty). Using the optimal hyperparameters, predictive probabilities were computed for all genes.

For the results in Section 2.5, for each species the pglm algorithm was retrained on the full GenAge dataset using GO features only. This choice of feature set was used to enable interpretation of regression coefficients. Here, the hyperparameter grid was the same 21 alpha values and 97 automatically selected lambda values with five-fold CV. The optimal alpha values were 0.15 for worm and 0.10 for yeast (both closer to ridge than lasso).

To facilitate reproducibility, all code used to produce the results in this manuscript is publicly available at https://github.com/willtownes/longevity-paper (Townes, Carr, & Miller, 2020).

## Acknowledgements

FWT was supported by NIH grant T32CA009337. KC was supported by the Vasilios Stavros Lagakos Fellowship. JWM gratefully acknowledges support from the Harvard Data Science Initiative Competitive Research Fund and the McLennan Family Fund. The authors thank Sheila Gaynor for advice on initial data cleaning, and Will Mair for helpful suggestions on biological interpretations. The original authors of McCormick et al. (2015) and Kemmeren et al. (2014) provided helpful advice in understanding their data, for which the authors are grateful. The authors declare no conflicts of interest.

## Author Contributions

JWM and KC conceived of the study. FWT and KC assembled and cleaned the datasets. KC performed initial analyses. FWT performed additional analyses, and drafted the initial report under JWM’s supervision. JWM revised the manuscript and provided biological interpretations of results. All authors approved of the final manuscript.

## Supplemental Figures and Tables

**Table S1:**
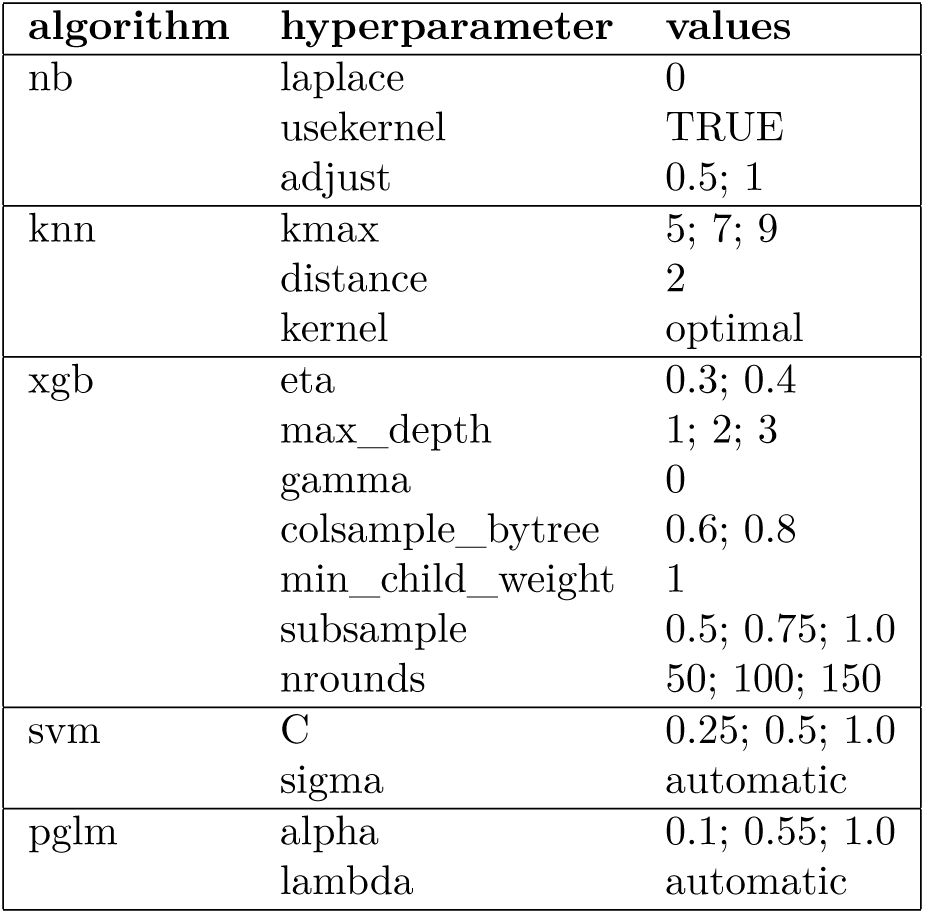
Hyperparameters used. pglm: elastic net penalized logistic regression, svm: support vector machine with radial basis function, xgb: gradient boosted trees, nb: naive Bayes, knn: k-nearest neighbors.

**Table S2:** Top worm pro-longevity genes not in GenAge predicted only using GO terms as features. Compare to Table 1.

| Gene | Prob | ID | Description |
| --- | --- | --- | --- |
| HLH-2 | 0.942 | WBGene00001949 | Helix Loop Helix |
| NPP-20 | 0.939 | WBGene00003806 | Protein SEC13 homolog |
| HMP-2 | 0.916 | WBGene00001979 | Beta-catenin-like protein hmp-2 |
| KGB-1 | 0.909 | WBGene00002187 | GLH-binding kinase 1 |
| NPP-13 | 0.908 | WBGene00003799 | Nuclear pore protein |
| DCS-1 | 0.905 | WBGene00000940 | m7GpppX diphosphatase |
| APR-1 | 0.903 | WBGene00000156 | Adenomatous polyposis coli protein-related protein 1 |
| UNC-51 | 0.892 | WBGene00006786 | Serine/threonine-protein kinase unc-51 |
| EMO-1 | 0.891 | WBGene00001303 | Protein transport protein Sec61 subunit gamma |
| PAS-1 | 0.891 | WBGene00003922 | Proteasome subunit alpha type-6 |

**Table S3:**
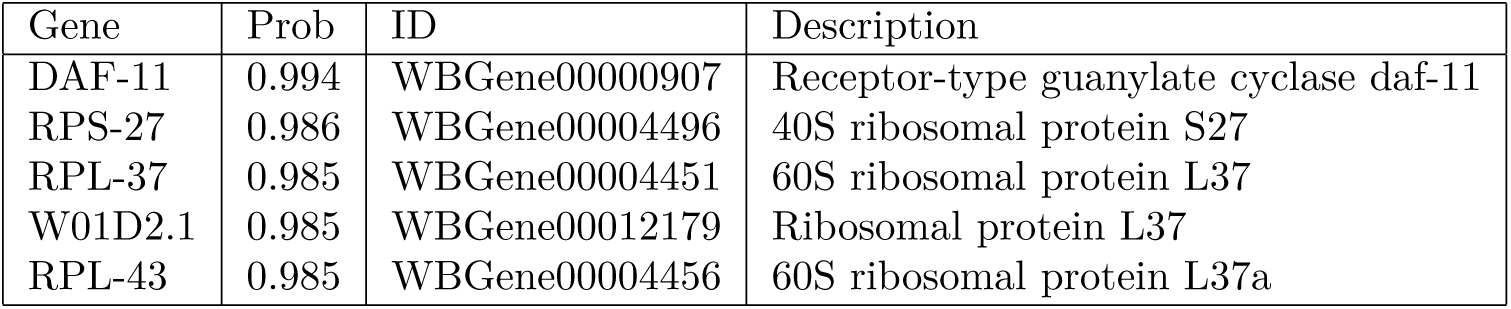
Top worm anti-longevity genes not in GenAge predicted only using GO terms as features. Compare to Table 1.

**Table S4:**
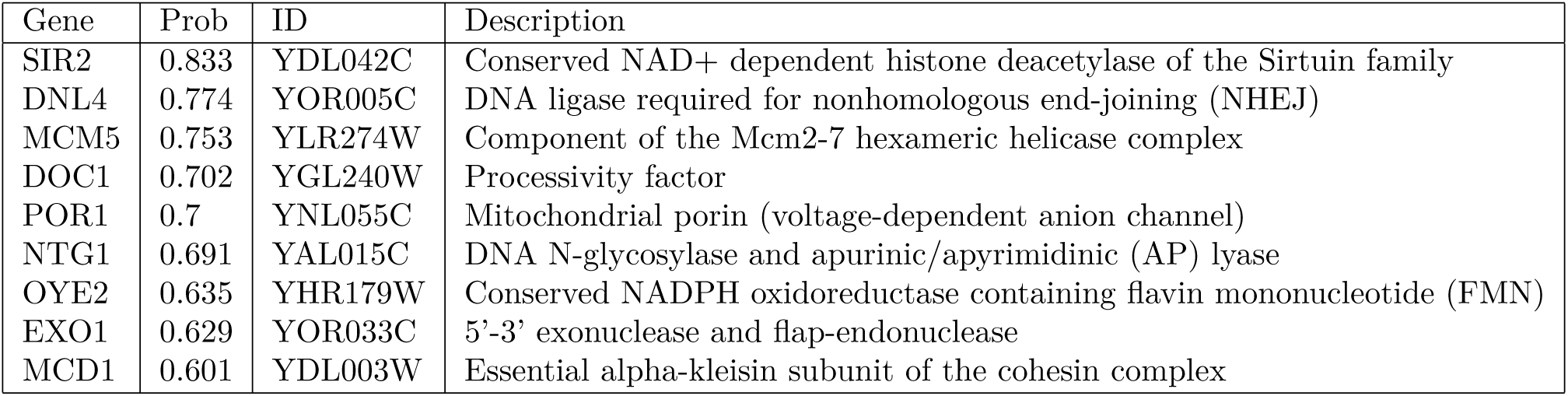
Top yeast pro-longevity genes not in GenAge predicted only using GO terms as features. Compare to Table 1.

**Table S5:**
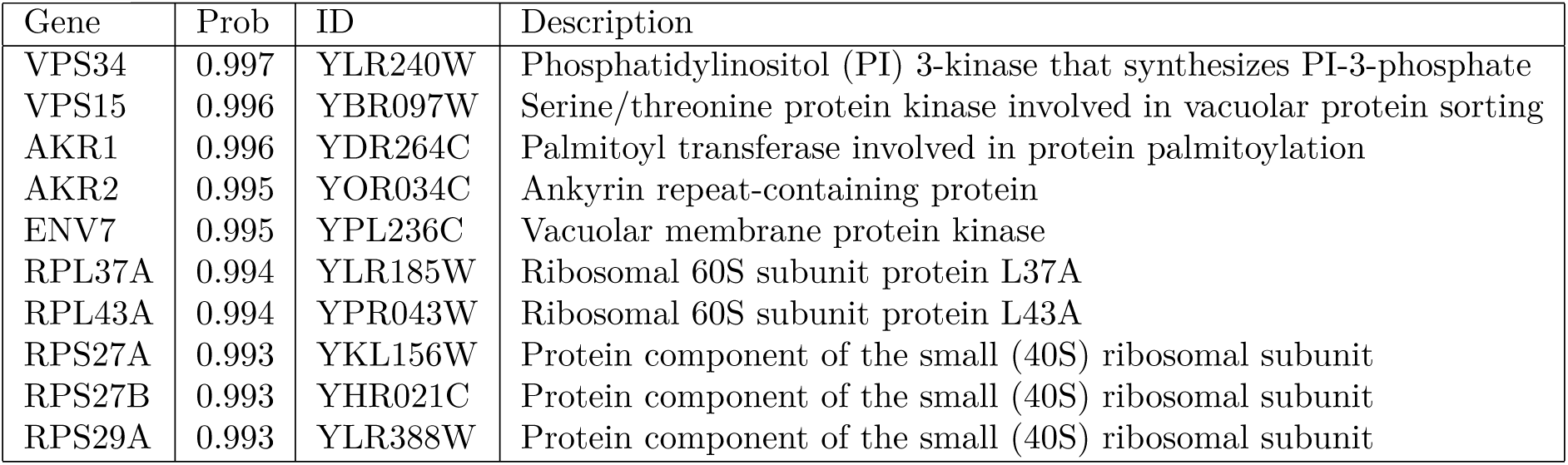
Top yeast anti-longevity genes not in GenAge predicted only using GO terms as features. Compare to Table 1.

**Figure S1:**
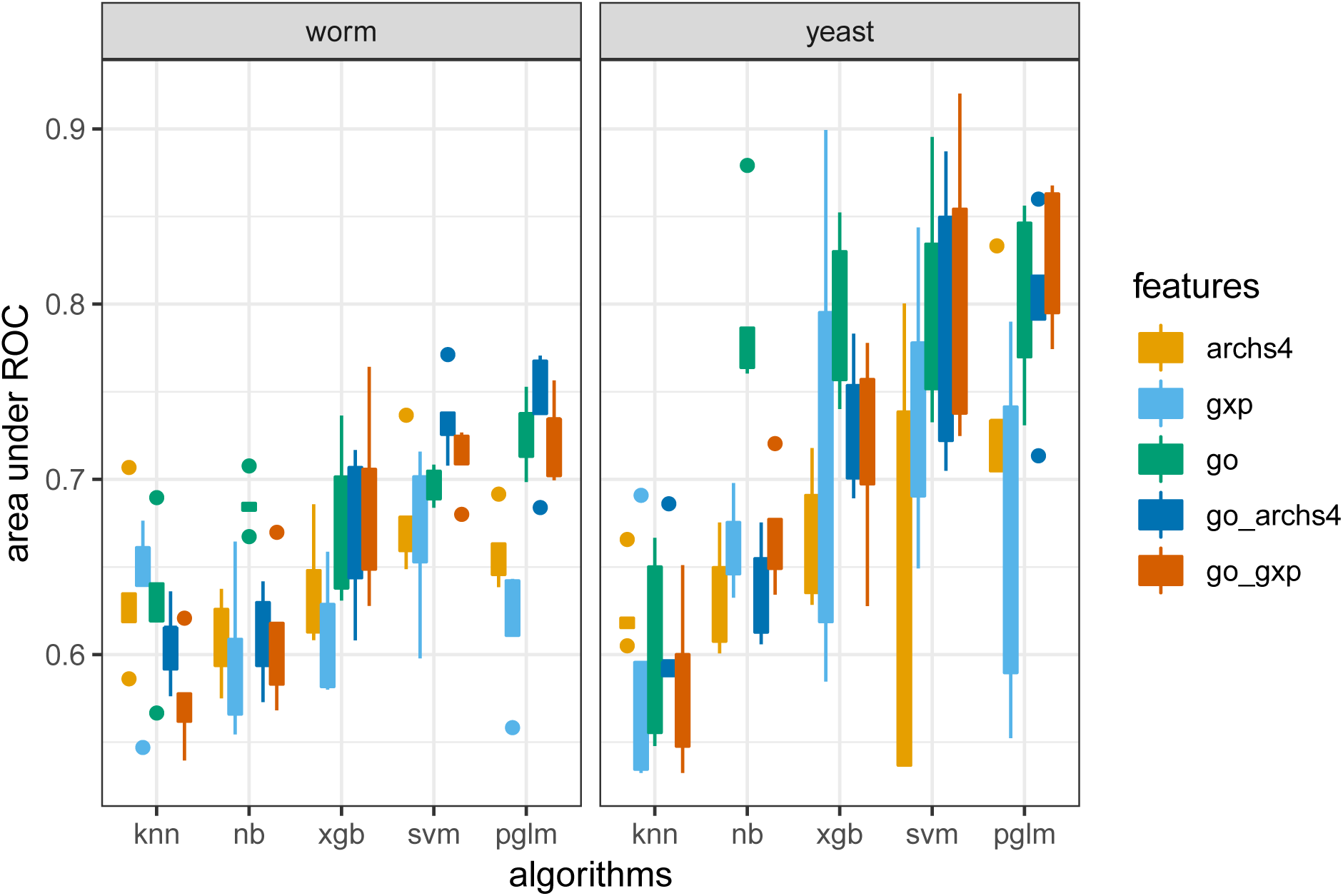
Comparison of predictive performance of machine learning algorithms on classifying genes as pro- or anti-longevity. pglm: elastic net penalized logistic regression, svm: support vector machine with radial basis function, xgb: gradient boosted trees, nb: naive Bayes, knn: k-nearest neighbors, gxp: gene expression, ROC: receiver-operator curve.

**Figure S2:**
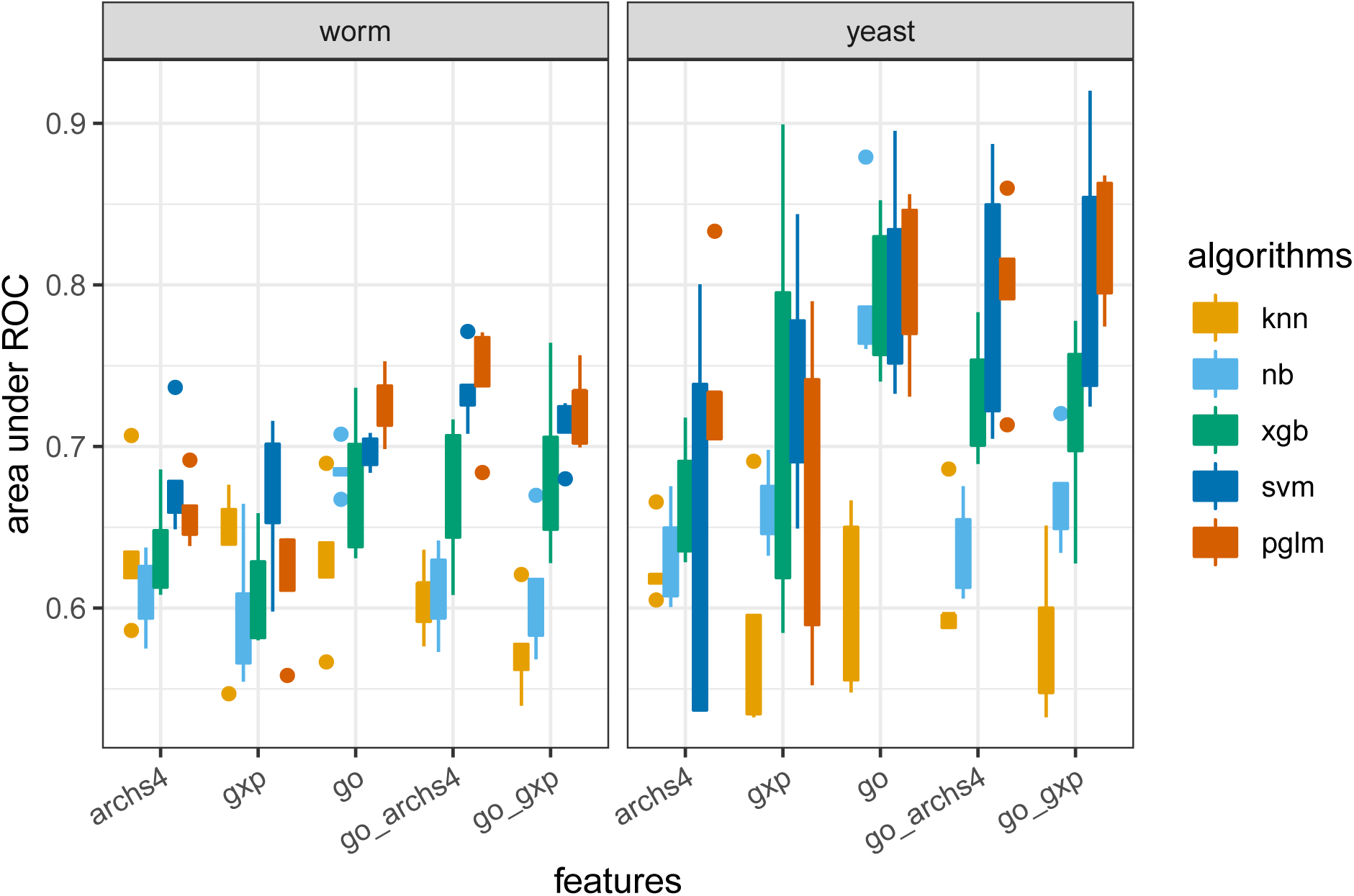
Comparison of predictive performance of different feature sets on classifying genes as pro- or anti-longevity. pglm: elastic net penalized logistic regression, svm: support vector machine with radial basis function, xgb: gradient boosted trees, nb: naive Bayes, knn: k-nearest neighbors, gxp: gene expression, ROC: receiver-operator curve.

**Figure S3:**
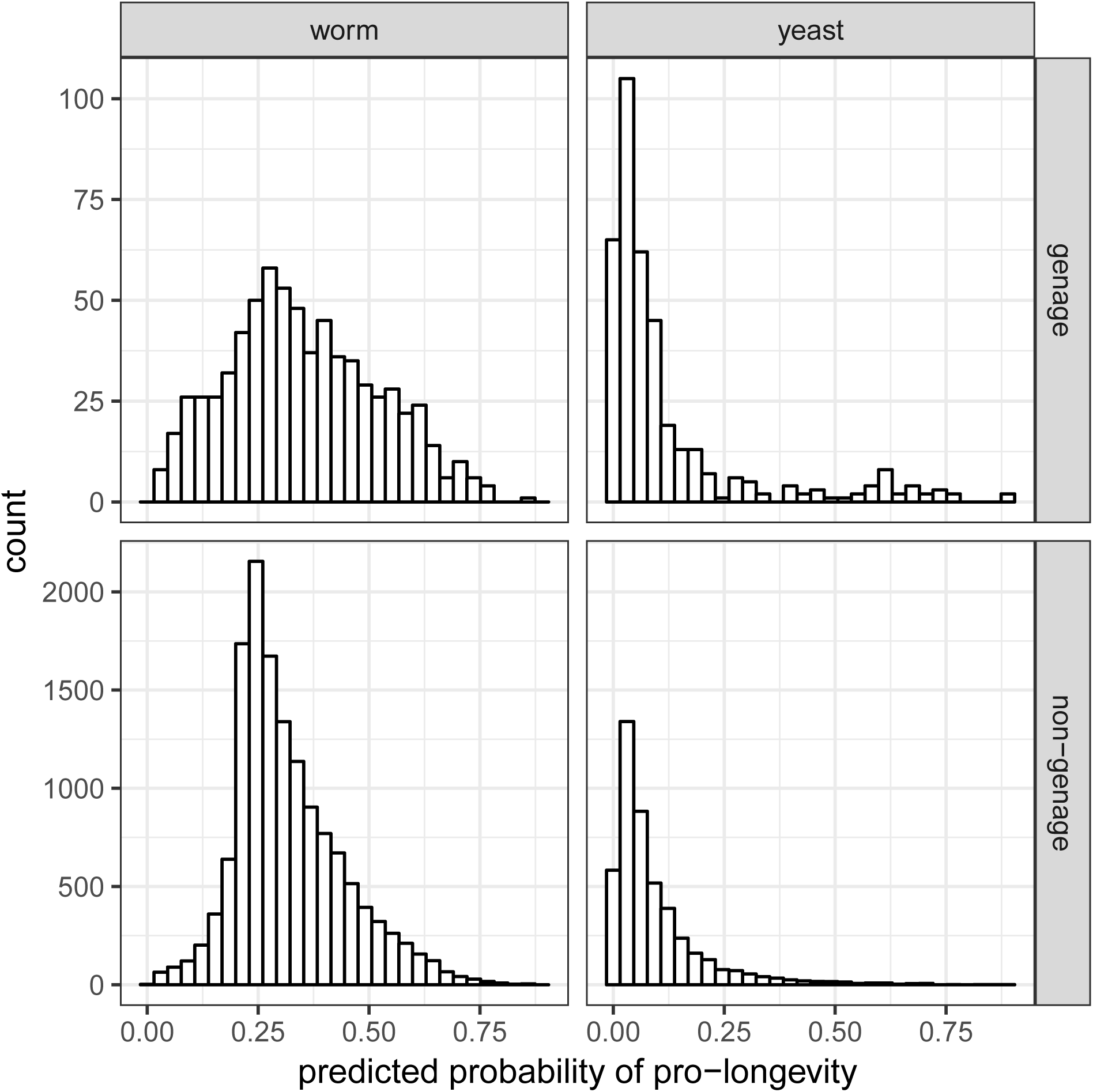
Distribution of predictive probabilities after training elastic net penalized logistic regression (pglm) on the full GenAge dataset with GO terms and ARCHS4 gene expression as features.

## References

Ailion, M., Inoue, T., Weaver, C. I., Holdcraft, R. W., & Thomas, J. H. (1999, June). Neurosecretory control of aging in Caenorhabditis elegans. Proceedings of the National Academy of Sciences of the United States of America, 96(13), 7394–7397.

Austriaco, N. R., & Guarente, L. P. (1997). Changes of telomere length cause reciprocal changes in the lifespan of mother cells in Saccharomyces cerevisiae. Proceedings of the National Academy of Sciences, 94(18), 9768–9772.

Ayyadevara, S., Dandapat, A., Singh, S. P., Siegel, E. R., Reis, R. J. S., Zimniak, L., & Zimniak, P. (2007). Life span and stress resistance of Caenorhabditis elegans are differentially affected by glutathione transferases metabolizing 4-hydroxynon-2-enal. Mechanisms of Ageing and Development, 128(2), 196–205.

Ayyadevara, S., Engle, M. R., Singh, S. P., Dandapat, A., Lichti, C. F., Beneš, H., … Zimniak, P. (2005). Lifespan and stress resistance of Caenorhabditis elegans are increased by expression of glutathione transferases capable of metabolizing the lipid peroxidation product 4-hydroxynonenal. Aging cell, 4(5), 257–271.

B Hwang, A., Jeong, D.-E., & Lee, S.-J. (2012). Mitochondria and organismal longevity. Current Genomics, 13(7), 519–532.

Cao, J., Packer, J. S., Ramani, V., Cusanovich, D. A., Huynh, C., Daza, R., … Shendure, J. (2017, August). Comprehensive single-cell transcriptional profiling of a multicellular organism. Science, 357 (6352), 661–667. doi:10.1126/science.aam8940

Carmona-Gutierrez, D., Hughes, A. L., Madeo, F., & Ruckenstuhl, C. (2016). The crucial impact of lysosomes in aging and longevity. Ageing Research Reviews, 32, 2–12.

Chen, L., Zhang, J., Xu, J., Wan, L., Teng, K., Xiang, J., … Liu, Xin (2016). rBm*α*TX14 increases the life span and promotes the locomotion of Caenorhabditis elegans. PLOS One, 11(9), e0161847.

Chen, T., He, T., Benesty, M., Khotilovich, V., Tang, Y., Cho, H., … Li, Y. (2019). Xgboost: Extreme Gradient Boosting.

Chen, X., McCue, H. V., Wong, S. Q., Kashyap, S. S., Kraemer, B. C., Barclay, J. W., … Morgan, A. (2015). Ethosuximide ameliorates neurodegenerative disease phenotypes by modulating DAF-16/FOXO target gene expression. Molecular Neurodegeneration, 10(1), 51.

Durieux, J., Wolff, S., & Dillin, A. (2011). The cell-non-autonomous nature of electron transport chain-mediated longevity. Cell, 144(1), 79–91.

Fabris, F., de Magalhães, J. P., & Freitas, A. A. (2017, April). A review of supervised machine learning applied to ageing research. Biogerontology, 18(2), 171–188. doi:10.1007/s10522-017-9683-y

Fang, Y., Fliss, A. E., Rao, J., & Caplan, A. J. (1998). SBA1 encodes a yeast hsp90 cochaperone that is homologous to vertebrate p23 proteins. Molecular and Cellular Biology, 18(7), 3727–3734.

Friedman, J., Hastie, T., & Tibshirani, R. (2010). Regularization Paths for Generalized Linear Models via Coordinate Descent. Journal of Statistical Software, 33(1), 1–22.

Garay, E., Campos, S. E., de la Cruz, J. G., Gaspar, A. P., Jinich, A., & DeLuna, A. (2014, February). High-Resolution Profiling of Stationary-Phase Survival Reveals Yeast Longevity Factors and Their Genetic Interactions. PLOS Genetics, 10(2), e1004168. doi:10.1371/journal.pgen.1004168

Gelman, A. (2008, July). Scaling regression inputs by dividing by two standard deviations. Statistics in Medicine, 27 (15), 2865–2873. doi:10.1002/sim.3107

Gene Ontology Consortium. (2019, January). The Gene Ontology Resource: 20 years and still GOing strong. Nucleic Acids Research, 47 (D1), D330–D338. doi:10.1093/nar/gky1055

Halaschek-Wiener, J., Khattra, J. S., McKay, S., Pouzyrev, A., Stott, J. M., Yang, G. S., … Riddle, Donald L (2005). Analysis of long-lived C. elegans daf-2 mutants using serial analysis of gene expression. Genome Research, 15(5), 603–615.

Hansen, M., Taubert, S., Crawford, D., Libina, N., Lee, S.-J., & Kenyon, C. (2007). Lifespan extension by conditions that inhibit translation in Caenorhabditis elegans. Aging Cell, 6(1), 95–110.

Haynes, W. A., Tomczak, A., & Khatri, P. (2018, January). Gene annotation bias impedes biomedical research. Scientific Reports, 8(1), 1–7. doi:10.1038/s41598-018-19333-x

Herbert, A. P., Riesen, M., Bloxam, L., Kosmidou, E., Wareing, B. M., Johnson, J. R., … Morgan, A. (2012). NMR structure of Hsp12, a protein induced by and required for dietary restriction-induced lifespan extension in yeast. PLOS One, 7 (7), e41975.

Hyun, M., Kim, J., Dumur, C., Schroeder, F. C., & You, Y.-J. (2016). BLIMP-1/BLMP-1 and metastasis-associated protein regulate stress resistant development in Caenorhabditis elegans. Genetics, 203(4), 1721–1732.

Jazwinski, S. M. (2005). Yeast longevity and aging—the mitochondrial connection. Mechanisms of Ageing and Development, 126(2), 243–248.

Johnson, T. E., & Lithgow, G. J. (1992). The Search for the Genetic Basis of Aging: The Identification of Gerontogenes in the Nematode Caenorhabditis elegans. Journal of the American Geriatrics Society, 40(9), 936–945. doi:10.1111/j.1532-5415.1992.tb01993.x

Karatzoglou, A., Smola, A., Hornik, K., & Zeileis, A. (2004). Kernlab – An S4 Package for Kernel Methods in R. Journal of Statistical Software, 11(9), 1–20.

Kemmeren, P., Sameith, K., van de Pasch, L. A. L., Benschop, J. J., Lenstra, T. L., Margaritis, T., … Holstege, F. C. P. (2014, April). Large-Scale Genetic Perturbations Reveal Regulatory Networks and an Abundance of Gene-Specific Repressors. Cell, 157 (3), 740–752. doi:10.1016/j.cell.2014.02.054

Kimura, K., Tanaka, N., Nakamura, N., Takano, S., & Ohkuma, S. (2007). Knockdown of mitochondrial heat shock protein 70 promotes progeria-like phenotypes in Caenorhabditis elegans. Journal of Biological Chemistry, 282(8), 5910–5918.

Kruegel, U., Robison, B., Dange, T., Kahlert, G., Delaney, J. R., Kotireddy, S., … Schmidt, Marion (2011). Elevated proteasome capacity extends replicative lifespan in Saccharomyces cerevisiae. PLOS Genetics, 7 (9), e1002253.

Kuhn, M., Weston, S., Williams, A., Keefer, C., Engelhardt, A., Cooper, T., … Hunt, T. (2019). Caret: Classification and Regression Training.

Lachmann, A., Torre, D., Keenan, A. B., Jagodnik, K. M., Lee, H. J., Wang, L., … Ma’ayan, A. (2018, April). Massive mining of publicly available RNA-seq data from human and mouse. Nature Communications, 9(1), 1366. doi:10.1038/s41467-018-03751-6

Lans, H., Lindvall, J., Thijssen, K., Karambelas, A., Cupac, D., Fensgård, Ø., … Vermeulen, W. (2013). DNA damage leads to progressive replicative decline but extends the life span of long-lived mutant animals. Cell Death and Differentiation, 20(12), 1709.

Liu, J., Wang, L., Wang, Z., & Liu, J.-P. (2019). Roles of telomere biology in cell senescence, replicative and chronological ageing. Cells, 8(1), 54.

Majka, M. (2019). Naivebayes: High Performance Implementation of the Naive Bayes Algorithm.

Marek, A., & Korona, R. (2013). Restricted Pleiotropy Facilitates Mutational Erosion of Major Life-History Traits. Evolution, 67 (11), 3077–3086. doi:10.1111/evo.12196

McCormick, M. A., Delaney, J. R., Tsuchiya, M., Tsuchiyama, S., Shemorry, A., Sim, S., … Kennedy, B. K. (2015, November). A Comprehensive Analysis of Replicative Lifespan in 4,698 Single-Gene Deletion Strains Uncovers Conserved Mechanisms of Aging. Cell Metabolism, 22(5), 895–906. doi:10.1016/j.cmet.2015.09.008

Narayan, V., Ly, T., Pourkarimi, E., Murillo, A. B., Gartner, A., Lamond, A. I., & Kenyon, C. (2016). Deep proteome analysis identifies age-related processes in C. elegans. Cell Systems, 3(2), 144–159.

Patil, C., & Walter, P. (2001). Intracellular signaling from the endoplasmic reticulum to the nucleus: The unfolded protein response in yeast and mammals. Current Opinion in Cell Biology, 13(3), 349–355.

Postnikoff, S. D., Johnson, J. E., & Tyler, J. K. (2017). The integrated stress response in budding yeast lifespan extension. Microbial Cell, 4(11), 368.

Remolina, S. C., Chang, P. L., Leips, J., Nuzhdin, S. V., & Hughes, K. A. (2012). Genomic Basis of Aging and Life-History Evolution in Drosophila Melanogaster. Evolution, 66(11), 3390–3403. doi:10.1111/j.1558-5646.2012.01710.x

Sampaio-Marques, B., & Ludovico, P. (2018). Linking cellular proteostasis to yeast longevity. FEMS Yeast Research, 18(5), foy043.

Schleit, J., Johnson, S. C., Bennett, C. F., Simko, M., Trongtham, N., Castanza, A., … Kaeberlein, M. (2013). Molecular mechanisms underlying genotype-dependent responses to dietary restriction. Aging Cell, 12(6), 1050–1061. doi:10.1111/acel.12130

Schliep, K., & Hechenbichler, K. (2016). Kknn: Weighted k-Nearest Neighbors.

Seufert, W., & Jentsch, S. (1990). Ubiquitin-conjugating enzymes UBC4 and UBC5 mediate selective degradation of short-lived and abnormal proteins. The EMBO Journal, 9(2), 543–550.

Shimasaki, T., Ohtsuka, H., Naito, C., Azuma, K., Tenno, T., Hiroaki, H., … Aiba, H. (2017). Ecl1 is a zinc-binding protein involved in the zinc-limitation-dependent extension of chronological life span in fission yeast. Molecular Genetics and Genomics, 292(2), 475–481.

Steffen, K. K., MacKay, V. L., Kerr, E. O., Tsuchiya, M., Hu, D., Fox, L. A., … Kaeberlein, Matt (2008). Yeast life span extension by depletion of 60s ribosomal subunits is mediated by Gcn4. Cell, 133(2), 292–302.

Sun, N., Youle, R. J., & Finkel, T. (2016). The mitochondrial basis of aging. Molecular Cell, 61(5), 654–666.

Sural, S., Lu, T.-C., Jung, S. A., & Hsu, A.-L. (2019). HSB-1 inhibition and HSF-1 overexpression trigger overlapping transcriptional changes to promote longevity in Caenorhabditis elegans. G3: Genes,Genomes, Genetics, 9(5), 1679–1692.

Tacutu, R., Thornton, D., Johnson, E., Budovsky, A., Barardo, D., Craig, T., … de Magalhães, J. P. (2018, January). Human Ageing Genomic Resources: New and updated databases. Nucleic Acids Research, 46(D1), D1083–D1090. doi:10.1093/nar/gkx1042

Tihanyi, B., Vellai, T., Regős, Á., Ari, E., Müller, F., & Takács-Vellai, K. (2010). The C. elegans Hox gene ceh-13 regulates cell migration and fusion in a non-colinear way. Implications for the early evolution of Hox clusters. BMC Developmental Biology, 10(1), 78.

Toogun, O. A., Zeiger, W., & Freeman, B. C. (2007). The p23 molecular chaperone promotes functional telomerase complexes through DNA dissociation. Proceedings of the National Academy of Sciences, 104(14), 5765–5770.

Townes, F. W., Carr, K., & Miller, J. W. (2020). Longevity paper github repository. https://github.com/willtownes/longevity-paper.

Van Voorhies, W. A. (1992). Production of sperm reduces nematode lifespan. Nature, 360(6403), 456.

Wilkinson, M. F. (2019, April). Genetic paradox explained by nonsense. Nature, 568(7751), 179. doi:10.1038/d41586-019-00823-5

Wu, L., Zhou, B., Oshiro-Rapley, N., Li, M., Paulo, J. A., Webster, C. M., … Soukas, Alexander A (2016). An ancient, unified mechanism for metformin growth inhibition in C. elegans and cancer. Cell, 167 (7), 1705–1718.

Yokoyama, K., Fukumoto, K., Murakami, T., Harada, S.-i., Hosono, R., Wadhwa, R., … Ohkuma, S. (2002). Extended longevity of Caenorhabditis elegans by knocking in extra copies of hsp70F, a homolog of mot-2 (mortalin)/mthsp70/Grp75. FEBS Letters, 516(1-3), 53–57.

